# Ideal free distribution of unequal competitors: spatial assortment and evolutionary diversification of competitive ability

**DOI:** 10.1101/2022.06.27.497704

**Authors:** Christoph Netz, Aparajitha Ramesh, Franz J. Weissing

**Affiliations:** Groningen Institute for Evolutionary Life Sciences, University of Groningen, Groningen, The Netherlands

**Keywords:** resource competition, patch choice, environmental change, evolutionary branching, individual differences, animal personality

## Abstract

Ideal free distribution theory attempts to predict the distribution of well-informed (‘ideal’) and unconstrained (‘free’) foragers in space based on adaptive individual decisions. When individuals differ in competitive ability, a whole array of equilibrium distributions is possible, and it is unclear which of these distributions are most likely. In the first part of our study, we show that strong competitors have an intrinsically stronger preference for highly productive habitat patches than poor competitors. This leads to an equilibrium distribution where the average competitive ability on a patch is strongly correlated with the productivity of the patch. In the second part of our study, we consider what happens if differences in competitive ability are heritable and, hence, subject to natural selection. Under constant environmental conditions, selection eliminates such differences: a single strategy prevails that optimally balances the costs and benefits associated with competitive ability. If the productivity of patches changes during the lifetime of individuals, the spatial assortment of competitors of equal competitive ability gives poor competitors a systematic advantage in times of environmental change, while good competitors benefit from equilibrium conditions. Using evolutionary individual-based simulations, we demonstrate that environmental change may then lead to the diversification of competitive ability.

## Introduction

Animals constantly have to make decisions on movement within or between habitats, especially in variable environments. The distribution of individuals depends on these decisions, which take into account the properties of the habitat and the distribution of conspecifics. The simplest forager distribution model (Fretwell & Lucas 1970) assumes a population of identical foragers, which are ‘ideal’ in that they have complete knowledge of the distributions of resources and conspecifics and are ‘free’ in that they are unrestricted in their movement. Foragers are then expected to distribute such that any further movement between patches does not increase the intake of any individual, yielding the so-called ideal free distribution (IFD). If foragers do not interfere with each other and share resources equally, the distribution of foragers corresponds to the distribution of resources, termed ‘input matching’ (Parker 1978). Although the IFD serves as a useful null model, in reality, individuals are neither ‘ideal’ nor ‘free’, and there is increasing evidence that consistent individual differences influence habitat choice and spatial distributions (Ehlinger 1990, Holtmann et al. 2017, Bonnot et al. 2018, Schirmer et al. 2019, 2020). This development is both a challenge and an opportunity for the theoretical framework of the ideal free distribution.

Several models have studied the distribution of foragers by relaxing key assumptions of the IFD, for example considering individuals that behave idiosyncratically and in non-optimal ways (Jackson et al. 2004, Matsumura et al. 2010) or incorporating individual differences that affect optimal decision making (Holt & Barfield 2008, Edelaar et al. 2008), specific examples including body size (Price 1983, Railsback & Harvey 2002), gizzard size (van Gils et al. 2005) or competitive ability (Sutherland & Parker 1985, 1992, Houston & McNamara 1988, Van de Pol et al. 2007, Smallegange & Van der Meer 2009). In particular, individual variation in competitive ability has been the focus of several modelling studies. Such variation is incorporated into IFD models in two different ways. In interference competition models, competitive ability affects the impact of interference on individual intake rates (Sutherland & Parker 1992, Smallegange & Van der Meer 2009). In this case, IFD theory predicts the segregation of unequal competitors over resource patches, where the most competitive types accumulate on patches with the highest resource levels, while weaker competitors occur at the lower resource levels. In exploitation competition models, the competitive ability of an individual determines the individual’s share in the local resources, for example via the capacity to defend territories (Huxley 1934). In this case, IFD theory predicts that, at equilibrium, the competition intensity on each patch (= the sum of the competitive abilities of the occupants of the patch) is proportional to the resource abundance on that patch (Sutherland & Parker 1985, Sutherland & Parker 1992). Such an equilibrium distribution can be realized in many different ways, and in principle, it is possible that weak and strong competitors co-occur on all patches or that weak competitors accumulate on patches with the highest productivity. Sutherland and Parker (1985) hypothesised that the most likely distribution of foragers converges on the IFD with equal competitors, which corresponds to the situation where, at equilibrium, the distribution of competitive types is roughly the same for all occupied patches. In contrast, Houston and McNamara (1988) argued that strong competitors should be slightly overrepresented on resource-rich patches, simply as a consequence of the number of ways in which the equilibrium distribution can be realized. Further work showed that the sequence and mechanism, by which foragers distribute across both patches, can have a significant impact on the equilibrium distributions that are reached (Spencer et al. 1995, Houston & Lang 1998).

Virtually all theoretical work on the distribution of unequal competitors has only considered the choice between two patches. The first goal of this study is to extend the theory to a more fine-grained environment with multiple patches. In addition, we consider a whole spectrum of competitive abilities. We show that stronger and weaker competitors differ in their patch preferences and that stronger competitors have, in comparison to weaker competitors, a systematic bias in favour of resource-rich patches. One would therefore expect competitor assortment, where strong competitors accumulate on resource-rich patches, while weak competitors typically occur on resource-poor patches. By means of individual-based simulations, we will show that such assortment does indeed take place under exploitation competition and that the effect is much stronger than the ‘statistical mechanics’ approach of Houston and McNamara (1988) suggests.

Most studies on the distribution of unequal competitors assume that differences in competitive ability are fixed and externally given. In many situations, it is likely that such differences are at least partly heritable (Baldauf et al. 2014). This implies that competitive ability is an evolvable trait. Therefore, we can ask not only how individual variation in competitive ability influences habitat choice and spatial distributions but also how (variation in) competitive ability is shaped by natural selection. Addressing this question is the second goal of this study.

One might expect that natural selection has the tendency to eliminate all variation in competitive ability, thus leading to a single strategy that optimally balances the costs and benefits associated with a given level of competitive ability. With a simple argument and some evolutionary simulations, we will show that this is indeed the case if the environment is stable, that is, if the resource level per patch remains constant. Making use of the assortment result derived in the first part of our study, we then argue that the situation may be different in case of a changing environment. With a simulation study, we will demonstrate that, under changing conditions, selection can lead to the diversification of competitive ability.

Our twofold purpose is therefore to first investigate the equilibrium distributions emerging from individual-based patch choice decisions, and secondly to study the evolutionary dynamics that this scenario implicates. We present a) an analytical description of how habitat preferences depend on individuals’ competitive abilities, and b) a simulation model of how spatial assortment can lead to the diversification of competitive ability. We thus show that spatial distributions are not only determined by the interactions between unequal competitors but that the process of repeated redistribution can by itself propel the evolution of several competitive morphs.

## Models and Results

We consider a population distributed across a number of patches, each of which provides a constant influx of resources that is shared among the foragers present on the patch. This situation is commonly known as a ‘continuous input’ model (Tregenza 1995). Individuals differ in their competitive ability, that is, their ability to defend resource shares against competitors. The intake rate of an individual on a habitat patch with resource influx *R* depends on the relation of the individual’s competitive ability *c*_*i*_ to the ‘competition intensity’ *C* on this patch, which is defined as the sum of the competitive abilities of all individuals present. In line with earlier work (Houston and McNamara 1988, Sutherland & Parker 1992, Tregenza 1995), we assume that the individual can consume a fraction *c*_*i*_/*C* of the local resources, yielding the intake rate:

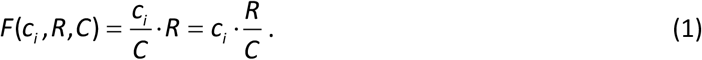

The ratio *R/C* may be viewed as the ‘resource availability’ on a given patch (per unit of competitive ability). As long as patches differ in their resource availability, at least some individuals have an incentive to move to a patch with higher resource availability. This will continue until an ‘ideal free distribution (IFD)’ is reached where all occupied patches have the same resource availability *R/C* (Sutherland & Parker 1985, Houston & McNamara 1988, Tregenza 1995).

### Spatial assortment: good competitors prefer resource-rich patches

At the ideal free distribution, the ratio *R/C* is equalized across all patches. Hence, the ideal free distribution depends on the distribution of competition intensity over patches and not directly on the distribution of individuals. In fact, many different distributions of foragers may lead to the same competition intensity on a given patch. For example, the same value *C* = 10 occurs when a patch is occupied by 10 individuals with competitive ability 1.0 or by 100 individuals with competitive ability 0.1. This implies that the IFD criterion (equality of the ratio *R/C*) can be satisfied by many different distributions of competitors over the patches. The question is whether some of these distributions are more likely than others. Sutherland and Parker (1985) predicted that the most likely distribution should correspond to the ideal free distribution with equal competitors since such a distribution corresponds to a random mixture of competitors over patches. Houston and McNamara (1988) noticed that among the many possible ways by which the IFD criterion can be satisfied those options where stronger competitors tend to occur on resource-rich patches are somewhat overrepresented. In analogy with statistical mechanics, they argue that it is, therefore, likely that at least some assortment of competitors over patches will occur. Although this argument is elegant, it is not immediately obvious whether principles of statistical mechanics can be applied to agents that do not move at random but by choosing the most suitable patch. Spencer et al. (1995) and Houston & Lang (1998) expanded on these results and showed that the sequence in which individuals move may have considerable influence on the resulting equilibrium distributions. Further, Houston & Lang showed that the movements of strong competitors may cause the subsequent movement of inferior competitors, providing a plausible mechanism by which spatial assortment may occur across patches. We here show that, more generally, the patch preferences of weaker competitors differ systematically from those of stronger competitors.

Consider an individual that compares two patches as to their suitability: patch 1 with resource influx *R*_1_ and current competitive intensity *C*_1_ and patch 2 with resource influx *R*_2_ and current competitive intensity *C*_2_. Assume further that patch 1 is the resource-richer patch: *R*_1_ > *R*_2_. An ideal and free individual with competitive ability *c*_*i*_ should prefer the resource-richer patch 1 if this patch, after the advent of the individual, yields a higher intake rate:

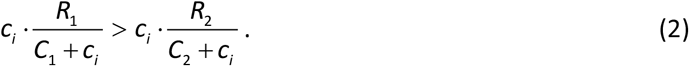

Notice that the denominators in (2) take account of the fact that the competition intensity of each patch would increase by *c*_*i*_, should our individual move to that patch. Inequality (2) is equivalent to:

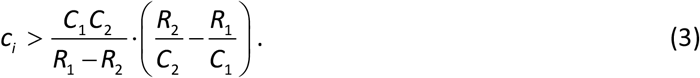

As long as the resource-rich patch 1 has a higher resource availability (*R*_1_/*C*_1_ > *R*_2_/*C*_2_), the right-hand side of (3) is negative, implying that *all* individuals prefer this patch, regardless of their competitive ability. This changes when the resource-rich patch 1 gets crowded to such an extent that the resource-poor patch 2 has a higher resource availability (*R*_2_/*C*_2_ > *R*_1_/*C*_1_). In this case, (3) is a threshold criterion: only those individuals with a sufficiently large competitive ability (larger than the right-hand side of (3)) will prefer the resource-rich patch 1, while individuals with lower competitive ability will prefer the resource-poor patch 2.

The above argument shows that individuals with a large competitive ability have a higher likelihood to prefer resource-rich patches than individuals with a smaller competitive ability. We therefore expect the assortment of competitive abilities along a resource gradient. To investigate the strength of this effect, we ran some individual-based simulations. We consider 100 patches with resource levels running from 0.01 to 1.0 at increments of 0.01. A population of 10,000 individuals containing the five different competitive types {0.1, 0.2, 0.4, 0.8, 1.6} in equal proportions is initially distributed randomly over the patches. Individual foragers are chosen in random order to compare intake rates among patches and move to the patch offering the highest intake rate. The individuals redistribute until no single individual can improve their intake rate any further, at which point a stable distribution is reached. As shown in Figure 1, the ensuing distributions are characterized by spatial assortment, where individuals of high competitive ability consistently occur more frequently on high resource patches, while individuals of low competitive ability occur on low resource patches. The degree of spatial assortment is surprisingly strong considering the relatively small influence of competitive ability on the comparison of potential intake rates between different patches (*C* >> *c*_*i*_, eqn (2)). As the IFD is approached, the difference between the *R/C* ratio of different patches becomes successively smaller, such that many patches offer relatively similar intake rates. In this case, the influence of individual competitive ability becomes temporarily decisive, producing the observed spatial correlations. As the differences between the *R/C* ratios decrease yet further, the threshold approaches zero and becomes irrelevant again.

**Figure 1:**
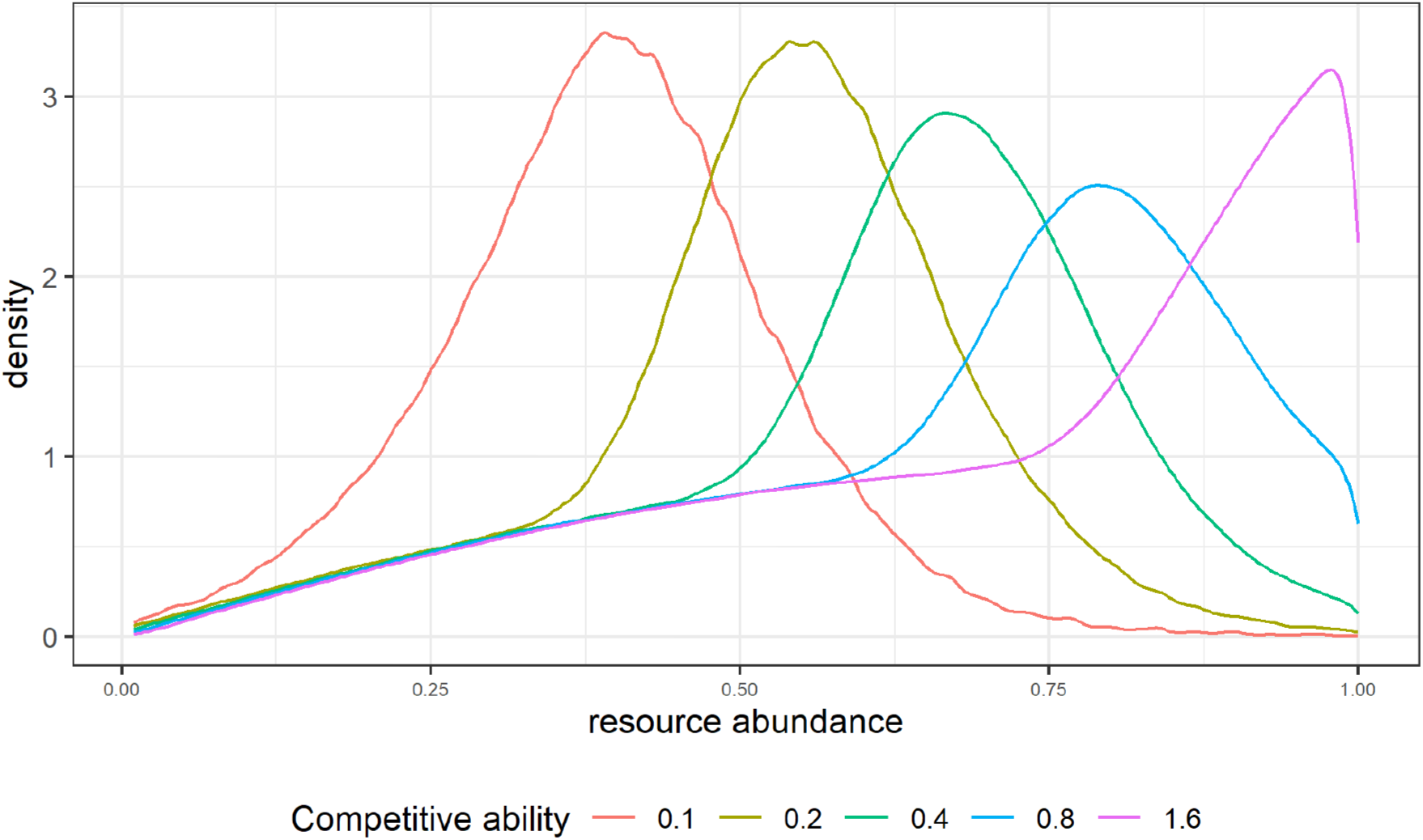
Ideal free distribution of unequal competitors over habitat patches differing in resource abundance. 10,000 individuals were initially distributed randomly over 100 patches with resource abundance values running from 0.01 to 1.00 at intervals of 0.01. One of five competitive ability values was randomly assigned to each individual. Then individuals moved sequentially (in random order) to the best-suited patch, until an ideal free distribution was reached. The graph shows the distribution of each competitive type at the IFD by combining the results of 100 replicate simulations.

### Evolution of competitive ability

Differences in individual competitive ability may arise at all levels from genetics to development and environmental effects during adulthood. From an evolutionary perspective, the presence of different types of competitors in a population poses the question of how multiple competitive types can coexist in a population. In the following we will consider how competitive abilities evolve in a patchy environment, first for a population that is permanently at the ideal free distribution (within generations) and second for a population where the IFD is repeatedly perturbed by changes in the environment.

In an evolutionary model, we have to specify how differences in intake rates translate into differences in survival and reproduction (Darwinian fitness). In optimal foraging models, either average food intake rate or lifetime resource consumption is typically taken as a proxy for fitness. When considering the evolution of competitive ability, this would not make much sense: according to eqn (1), the intake rate on each patch is proportional to an individual’s competitive ability. Hence, the highest possible competitive ability would evolve if it could be realized without costs. Here, we assume that a higher competitive ability is metabolically costly, and that the per-time-unit costs for a competitive ability *c*_*i*_ amount to *kc*_*i*_ resource units, where *k* is a constant of proportionality. Our fitness proxy is therefore based on the *net* intake rate:

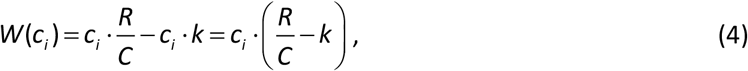

which, accumulated over the lifetime of an individual, is our measure of lifetime reproductive success. At the IFD, the resource availabilities *R*/*C* are equal across all patches and given by 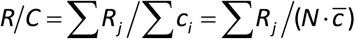, where *N* is the number of individuals and 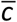 is their average competitive ability. If we insert this expression into (4), we can conclude that the net intake rate *W* increases with *c*_*i*_, if 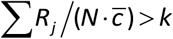 and decreases with *c*_*i*_, if 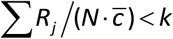. This implies that competitive ability will converge to a level *c*^*^ at which 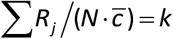. As the corresponding population is monomorphic, the value *c*^*^ is equal to the average competitive ability 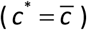. This yields:

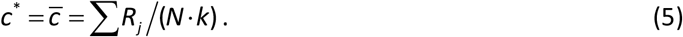

To check this expectation, we ran individual-based evolutionary simulations. Each individual is endowed with a heritable competitive ability. Within generations, individuals move to a patch yielding the maximal intake rate (given their competitive ability); movement will stop once the ideal free distribution is reached. Between generations, individuals produce offspring that inherit the competitive ability of their parent (subject to rare mutations). As the number of offspring is proportional to the net intake, accumulated over lifetime, those competitive abilities will increase in frequency that realize the highest net foraging success. A more detailed description of the model is provided in the appendix. Figure 2 shows that, irrespective of the initial conditions, the simulations evolve to the value of *c*^*^ predicted by eqn (5) and therefore confirm our analytical expectations.

**Figure 2:**
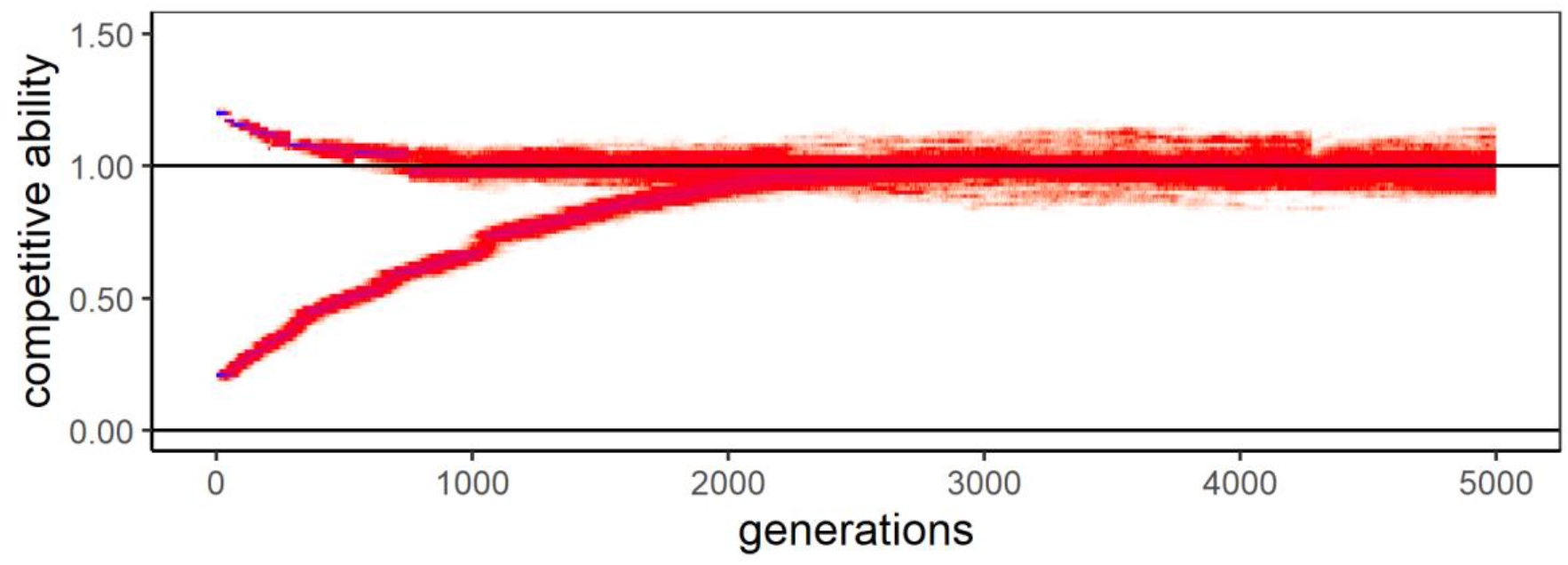
Evolution of competitive ability under IFD conditions. Two simulations, starting at different initial conditions, for the evolution of competitive ability in a system where 10,000 individuals distribute over 100 patches with resource abundances varying between 0 and 1. The cost parameter *k* had the value 0.005. Both simulations converge to the value *c*\* = 1.0, the value of competitive ability predicted by eqn (5). The relative frequencies of trait values within each generation are encoded by a colour gradient from 0.0 (= white) to 0.3 (= red) and 1.0 (= blue).

### Changing environments: evolution of competitive diversity

If environmental conditions remain constant within a generation, a population of foragers will rapidly converge to the IFD. Accordingly, the population will converge to a monomorphic state where all individuals have the same competitive ability *c*^*^. Some limited variation around *c*^*^ remains due to the ongoing influx of mutations (selection close to the evolutionary equilibrium is weak and not very efficient in eliminating mutations that are close to *c*^*^), but larger-scale variation in competitive ability is eliminated. Resource environments are rarely static, however, and the ideal free distribution is therefore often a fleeting target. If the environment changes repeatedly within a generation and if it takes time to re-establish the IFD after each change, it is no longer obvious that only a single competitive ability *c*^*^ will persist.

To see this, consider a population with variation in competitive abilities. As we have seen above, strong competitors will, under IFD conditions, accumulate on resource-rich patches, while weak competitors will mainly occur on resource-poor patches. If the environment (i.e., the resource influx per patch) changes at random, previously resource-rich patches will, on average, deteriorate while previously resource-poor patches will, on average, improve. This implies that changing conditions will, on average, be detrimental for strong competitors (that have accumulated on the previously resource-rich patches) and beneficial for weak competitors (that mainly occur on the previously resource-poor patches). It is conceivable that this principle will facilitate the coexistence of different competitive types, where in times of stasis (under IFD conditions), strong competitors have a higher net intake rate, while in times of change, weak competitors have a higher net intake rate.

To test this idea, we ran our evolutionary simulations under a stochastic regime of change, where the patch-specific resource levels changed at a rate of 0.25 (i.e., on average every 4 time units). In this variant of the model (see the appendix for details), foragers scan their environment at a rate of 0.5, thus noticing on average every 2 time units whether changes have occurred that may induce them to move to a patch with a higher net intake rate. Figure 3 shows that, under these changing conditions, evolution does indeed not lead to a monomorphic state. Instead, the population diversifies into a large number of coexisting competitive types.

**Figure 3:**
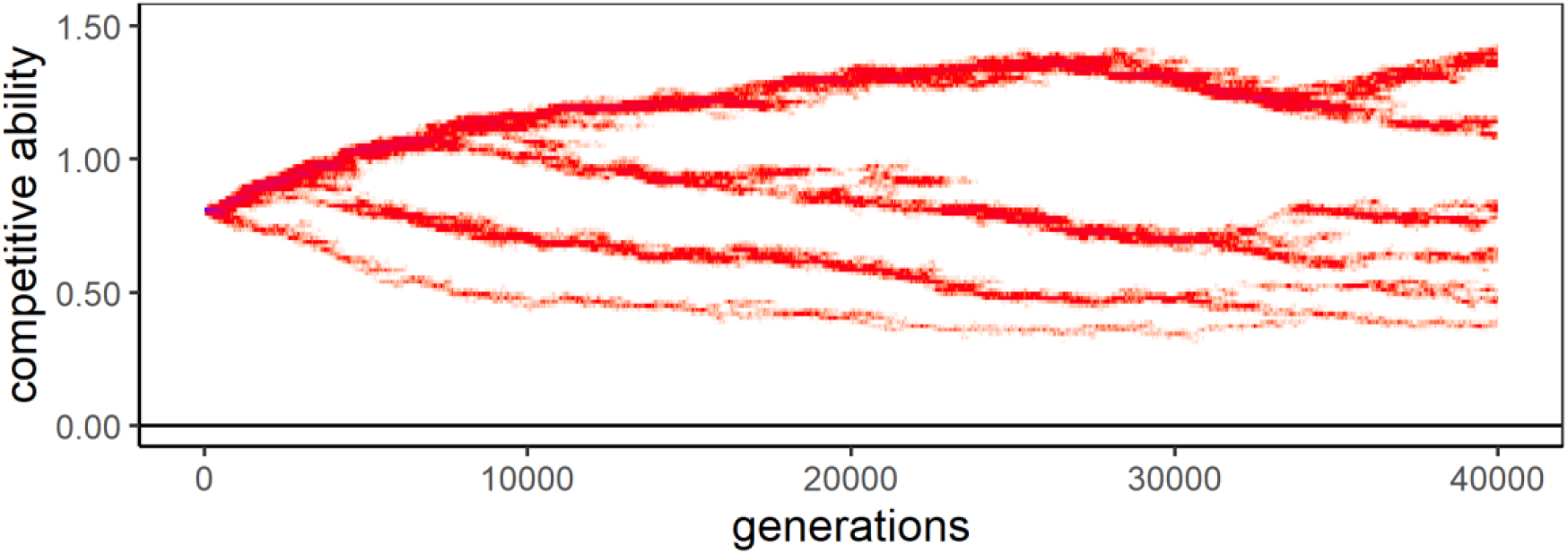
Evolutionary diversification of competitive abilities under changing environmental conditions. The graph shows one representative simulation for the same parameter settings as in Fig. 2. Now, however, the resource influx per habitat patch does not remain constant throughout a generation but randomly changes on average once every four time units. In the course of evolution, the population ‘branches’ into distinct competitive types.

Figure 4 demonstrates that, as predicted, the coexisting competitive types receive a differential net intake at equilibrium and after a change. Under stable conditions (when the population is close to the IFD), the net intake rates increases with competitive ability, while under changing conditions the weakest competitors have the highest net intake rate. The spatial assortment of less competitive individuals on poor patches and more competitive individuals on rich patches produces a transient benefit of spatiotemporal variation for the former.

**Figure 4:**
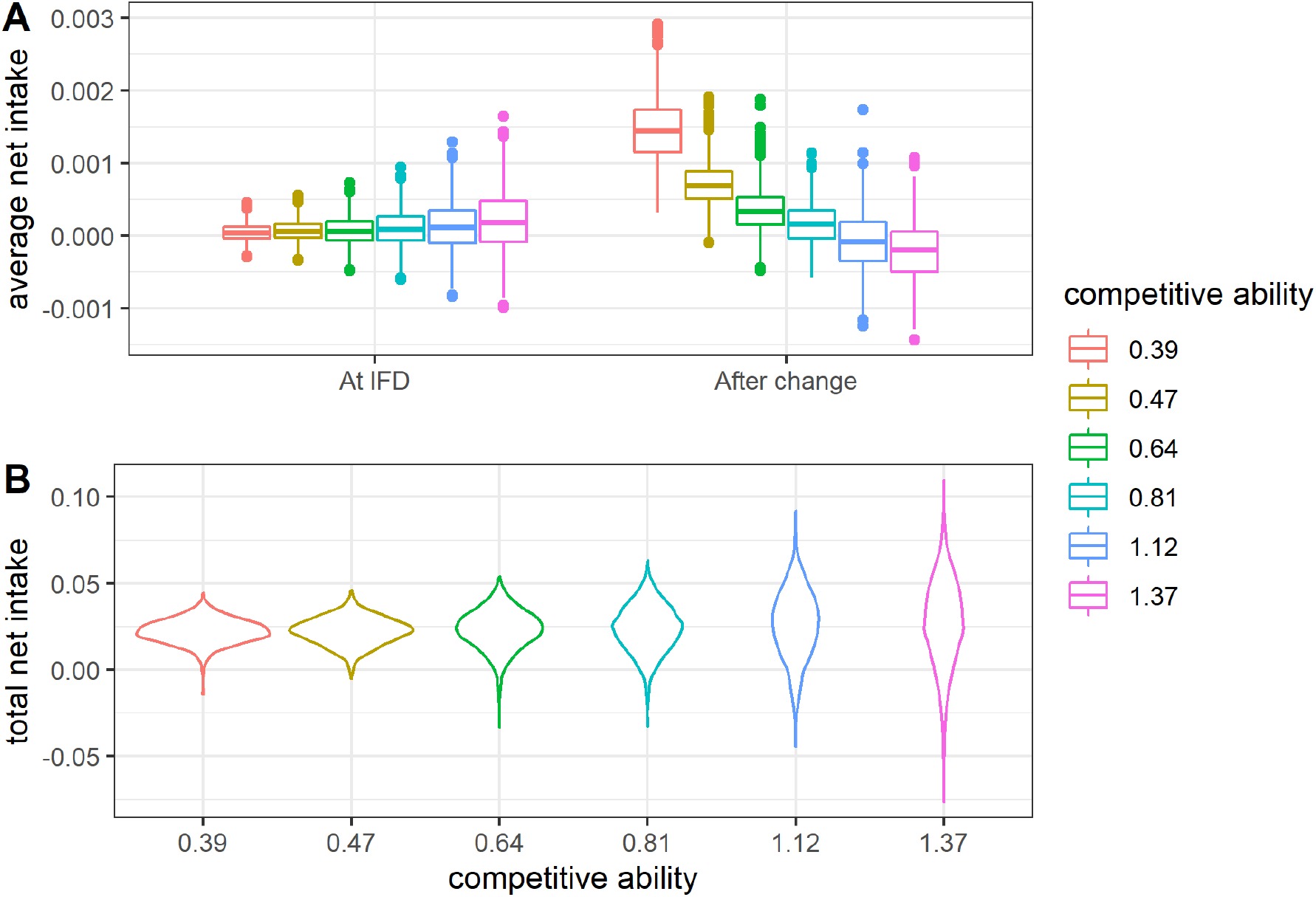
Net intake rates under changing environmental conditions. For the simulation in Fig. 3, we binned the six competitive types in generation 40,000 and (A) averaged their momentary net intake rates under IFD conditions (left part of the graph) and immediately after a change of the environment (right part of the graph). Net intake rate increases with competitive ability under stable conditions (at IFD), while it decreases with competitive ability under changing conditions. (B) The total net intake over individual lifetime is roughly the same for all six morphs.

## Discussion

Competition is a central motive in ecology and evolution and may determine forager distributions as well as the course of natural selection. We here considered the patch choice decisions of individuals, the equilibrium distributions emerging from these decisions, and the evolutionary dynamics of competitive abilities under stable and changing environmental conditions. We arrived at two key insights. First, the ranking of habitat patches as to their suitability (= net intake rate) is systematically affected by the competitive ability of the decision-making individual. Quite generally, strong competitors have a higher tendency to prefer resource-rich patches than weak competitors. Although this bias is relatively small, it can result in strong spatial assortment, where stronger competitors accumulate on resource-rich patches, while weaker competitors mainly occur on resource-poor patches. Second, this spatial assortment has important implications for the evolution of competitive ability. Under constant environmental conditions, variation in (heritable) competitive abilities cannot persist, and the population will converge to a monomorphic state with one type of competitor. If, however, environmental conditions change within generations, spatial assortment leads to a situation where strong competitors have an advantage under stable conditions (at IFD), while weak competitors have an advantage in periods of environmental change. As a consequence, foragers differing in competitive ability can have the same fitness (= net intake rate, summed or averaged over lifetime), allowing coexistence. We have shown that such polymorphism does indeed evolve: through repeated ‘evolutionary branching’ (Geritz et al. 1998, Baldauf et al. 2014), a large number of competitive types emerges and stably coexists.

In contrast to interference models, continuous input models, such as the one considered here, do not predict the segregation of unequal competitors, as the IFD condition (equality of resource abundance *R*/*C* across patches) can be satisfied in a multitude of ways. Sutherland & Parker (1985) and Parker & Sutherland (1986) speculated that unequal competitors will typically occur in roughly equal proportions at all patches, which would lead to the same IFD as predicted in the absence of differences in competitive ability. This is not the case in our model implementation, where at the IFD strong competitors are overrepresented on the resource-rich patches. For the special case of two patches, other studies (e.g., Houston & McNamara 1988, Spencer et al. 1995, Houston & Lang 1998) arrived at a similar conclusion, but based on different arguments. In Appendix B, we investigate in some detail how our findings relate to the results of these earlier studies. We confirm the findings of Spencer et al. (1995) and Houston & Lang (1998) that the degree of competitor assortment strongly depends on the way how individuals make their patch choice decisions, and we add one insight to those discussed in these papers. Both Spencer et al. (1995) and Houston & Lang (1998) consider foragers moving into the patches from the outside (a mechanism we call ‘external initialisation’): two initially empty patches fill up due to the sequential arrival of individuals, each newly arriving individual choosing the patch offering the highest intake rate. In contrast, our study considers an ‘internal initialisation’ scenario, where the individuals are initially distributed randomly over the patches and subsequently sequentially relocate themselves if another patch offers a higher intake rate. In case of two patches, we show (Fig. S1) that external initialisation leads to strong assortment, while internal initialisation does not lead to assortment at all. In other words, the distribution of ideal and free competitors over patches strongly depends on whether the competitors make their choices when entering the system from the outside (external initialisation) or from within (internal initialisation).

The no-assortment result of Fig. S1 points at an interesting discrepancy between the two-patch scenario typically considered in the literature and the multi-patch scenario considered in our study. Why does one of our key findings, assortment of competitors at a multi-patch IFD, break down for the special case of two patches? In Appendix B, we provide an explanation. We show that our threshold criterion (3) is generally (i.e., also for the case of two patches) applicable to the external initialisation scenario, and that it therefore explains the assortment results of Spencer et al. (1995) and Houston & Lang (1998). However, the criterion ceases to hold in the special case of two patches and internal initialisation, where it needs to be replaced by an alternative criterion (S2, see Appendix B), which no longer predicts assortment. Interestingly, assortment is re-established if the two patches are split into sub-patches that have the same properties as their ‘mother patch’ (Figure S2). This implies that the distribution of competitors over space may depend strongly on the ‘graininess’ of the environment. If, for example, the habitat choice situation is framed in a coarse-grained manner, such as a decision between deciduous and coniferous forest, our model would not predict assortment. In contrast, the same model would predict the accumulation of strong competitors in productive habitats if the otherwise identical situation is framed in a more fine-grained way, such as a decision between a multitude of deciduous and coniferous forest plots.

The existing models on the distribution of unequal competitors assume that differences in competitive ability are externally given. Such an analysis is incomplete if competitive differences have a heritable component. If this is the case, ideal free distribution theory, which is rooted in evolutionary optimality thinking (Netz et al. 2022), should pose the question whether unequal competitors can stably coexist in the course of evolution and, if so, how the distribution of competitive types is shaped by natural selection. We have shown that the evolutionary coexistence of unequal competitors is unlikely if the population is at an ideal free distribution all the time. This conclusion may change, however, if deviations from IFD conditions occur regularly. Such deviations are, for example, to be expected if sensory and/or locomotory constraints are taken into account (i.e., if the individuals are less ‘ideal’ and ‘free’ than IFD theory assumes). Here, we considered an alternative scenario, where IFD conditions are frequently perturbed due to environmental change. By means of a simple model, we demonstrated that distinct competitive types can emerge and stably coexist in the course of evolution. Consistent individual differences may therefore be as much a consequence as they are a cause of spatial distribution of individuals within the population (see also Wolf and Weissing 2010). As the evolved differences in phenotype (= competitive ability) lead to consistent differences in behavioural dispositions (= patch preferences; see (3)), we can conclude that spatiotemporal variation of the environment paves the way to the evolution of ‘personality’ differences.

It has been argued repeatedly (e.g., Dingemanse and Wolf 2010, Wolf and Weissing 2010, Dall et al. 2012) that spatiotemporal variation of the environment, coupled with constraints on matching the environment, may be a key driver of personality differences, but to our knowledge this has not been demonstrated in a formal model before. Empirical evidence for this is hard to collect in wild populations, but emergent spatial patterns have been studied in a number of taxa. In great tits, spatiotemporal variation in resources (here, nest boxes) within and between populations and study plots have been implicated in the coexistence of different exploratory tendencies (Nicolaus et al. 2016, Mouchet et al. 2021). Similarly, dispersal syndromes have been reported to be present in heterogeneous environments with fluctuations in habitat quality, risks and competition leading to spatial structuring of a population (Duckworth 2006, Cote et al. 2010), much like in our simulations. Taborsky et al. (2014) found that habitat competition between cichlids of different body sizes leads to assortment and ultimately assortative mating, which is another potent factor by which spatial distributions can affect the course of evolution in sexually reproducing species. There is also empirical evidence for habitat choice based on personality, leading to a biased spatial distribution of behavioural types and behaviour-environment correlations (Edelaar et al. 2008, Pearish et al. 2013, Holtmann et al. 2017). However, in these cases, the mechanisms underlying such spatial structuring of personality types are often in the dark.

Our two key results, the emergence of spatial assortment in a continuous input model of the IFD with unequal competitors, and the occurrence of polymorphism in an evolutionary model incorporating the very same, are both derived from an extension of a simple analytical model with certain mechanistic assumptions. We suggest that this is a constructive approach to study the robustness of these analytical models, and to uncover phenomena that would be otherwise overlooked. This model also acts as a useful starting point to relax further assumptions of IFD and extend to other dimensions of biologically relevant traits such as responsiveness to environmental change or limits to perception.

## Data and code availability

Simulation model and data analysis code is available on Github: https://github.com/christophnetz/unequal_competitors and as a stable release on Zenodo: https://doi.org/10.5281/zenodo.6634659

## Acknowledgements

The authors thank Jakob Gismann, Pratik Gupte and other members of the Modelling Adaptive Response Mechanisms Group at the University of Groningen for helpful discussions on the manuscript, as well as Hanno Hildenbrandt for the implementation of parallel threading. F.J.W. and C.N. acknowledge funding from the European Research Council (ERC Advanced Grant No. 789240).

## Appendix A. Description of the evolutionary simulation model

### Ecological setting

We consider 100 patches, with resource densities drawn from a uniform distribution between 0 and 1 at initialization and during every change of the environment.

Individual movements and environmental change occur in an event-based approach, where each event occurs at a constant rate. Individual foragers scan their environment at a rate of 0.5, compare the potential intake across all patches and move to the patch providing the highest intake rate. Environmental change occurs at a rate of 0.25, and therefore on average every 4 time units. For computational convenience, foragers consume resources at discrete intervals of one time unit.

### Reproduction and Inheritance

We consider discrete, non-overlapping generations of 100 time units, at the end of which reproduction occurs. For simplicity, reproduction is asexual. Individuals are haploid and have a single gene locus encoding for competitive ability that is inherited from parent to offspring. For each individual, the cumulative lifetime net intake *W*_*cum*_ is calculated. To prevent negative fitness values, a baseline value *W*_0_ is added to *W*_*cum*_, which can be interpreted as food intake that is unaffected by competitive interactions. The number of offspring produced per parent is determined by a weighted lottery that ensures that the expected number of offspring of an individual is proportional to *W*_*cum*_ +*W*_0_ and that population size remains constant at 10,000 individuals. Offspring inherit the competitive ability from their parent, subject to rare mutations of small effect size. Mutations occur at a rate of 0.01 per reproduction event. When a mutation occurs, a random number, drawn from a normal distribution with mean zero and standard deviation *σ* = 0.01, is added to the parental value. At the beginning of the new generation, offspring are randomly distributed over the patches.

## Appendix B. Comparison with two-patch models

For the special case of two habitat patches, Houston and McNamara (1988) showed that the distribution of competitors over patches at the IFD is biased in such a way that strong competitors are more likely to occur on the resource-rich patch. This result reflects the fact that among the many possible distributions satisfying the IFD condition, those with an accumulation of strong competitors on the resource-rich patch are overrepresented. To see this, consider two patches A and B, of which A is twice as resource-rich as B (*R*_*A*_ = 2*R*_*B*_). If all competitors are equal, 2/3 of all individuals would therefore occur in patch A in the ideal free distribution. Consider now two types of competitors, of which type 1 is twice as strong as type 2 (*c*_1_ = 2 *c*_2_); both types are equally frequent 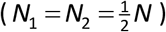. In Figure S1, the green curve shows the frequency distribution of the number of individuals in patch A for all realisations of the IFD condition. In the majority of cases, the number of individuals on patch A is smaller than 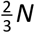, implying that the strong competitors are overrepresented on this resource-rich patch. The green distribution in Fig. S1 represents the complete set of IFD realisations, and the validity of Houston and McNamara’s ‘statistical mechanics’ argument relies on the assumption that the IFD that is actually realised is an unbiased sample of all IFD realisations.

**Figure S1:**
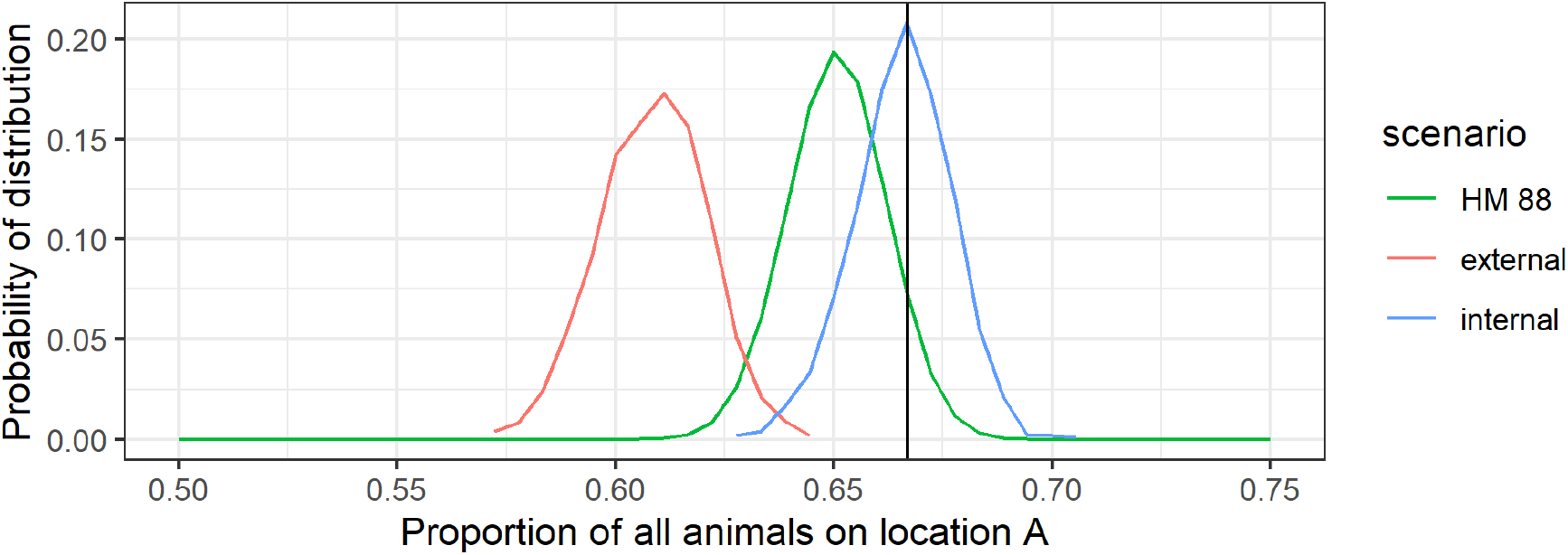
Implications of three habitat choice scenarios for the assortment of competitors. Following Houston & McNamara (1988), we consider a population of 180 individuals that distribute over two patches differing in quality. Resource abundance on patch A is twice the resource abundance on patch B. If all individuals were equal, 2/3 would occur on patch A at the IFD (vertical black line). Assume now that individuals differ in competitive ability: there are 90 good competitors that are twice as strong (*c*_1_ = 2 *c*_2_) as the 90 bad competitors. The <u>green curve</u> shows the probability distribution of the proportion of individuals on the resource-rich patch A, as derived from the ‘statistical mechanics’ analysis of Houston & McNamara (1988). The major part of this distribution is to the left of the value 2/3, indicating that, on average, strong competitors accumulate on the resource-rich patch. The <u>red curve</u> shows the probability distribution resulting from the ‘external initialisation’ scenario, where two initially empty patches fill up due to the sequential arrival of individuals, each newly arriving individual choosing the patch offering the highest intake rate. This choice scenario leads to an even stronger assortment of competitors to patches. The <u>blue curve</u> shows the probability distribution resulting from the ‘internal initialisation’ scenario, where the individuals are initially distributed randomly over the patches and subsequently sequentially relocate themselves if the other patch offers a higher intake rate. No assortment does occur in this scenario. The distributions shown are based on 1,000 replicate simulations per scenario.

A subsequent investigation by Houston and Lang (1998) showed that the distribution of actual IFD realisations strongly depends on the way the equilibrium distribution of competitors over patches is achieved. If, for example, the good competitors make their habitat choice decisions before the bad competitors, the number of individuals on the resource-rich patch will be 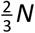 at the IFD, as in the case of equal competitors. If, in contrast, competitors make their decisions sequentially, in a random order, good competitors accumulate even more strongly on the resource-rich patch A than predicted by Houston & McNamara. In either case, the solution set calculated by Houston and McNamara (1988) is not representative for the realized distribution of competitors over patches.

An important detail of Houston and Lang’s (1988) treatment is that their individuals sequentially enter the two patches from the outside, whereas in our model we assume that the foragers are already distributed across the patches and subsequently redistribute until an IFD is reached. Figure S1 shows that the initialisation has a clear effect on the outcome. While ‘outside initialisation’ (red) leads to a pronounced assortment (i.e. the accumulation of strong competitors on the resource-rich patch A), this is not the case for the scenario where the individuals were first distributed randomly over the two patches (blue). In both cases, the realized distributions of competitors over patches are considerably different from the one predicted by Houston & McNamara (1988).

In view of our threshold criterion (inequality (3) in the main text), it is understandable that ‘outside initialisation’ leads to pronounced assortment: strong competitors have a higher tendency to choose the research-rich patch than weak competitors. But why does this argument break down in the case of ‘random initialisation’? We see two reasons for this. First, strong and weak competitors only differ in their patch preferences if the difference in resource availabilities (= the difference in *R*/*C*-values) is such that the right-hand side of (3) is larger than the lowest competitive ability *c*_min_ and smaller than the highest competitive ability *c*_max_. If the patches fill up sequentially (‘outside initialisation’), the resource availabilities *R*_*A*_/*C*_*A*_ and *R*_*B*_/*C*_*B*_ will, due to the choices of the newly arriving individuals, remain similar to each other, implying that the threshold criterion (3) will often lead to different outcomes for weak and strong competitors. If, in contrast, the patches are initialised at random, the resource availabilities will initially differ a lot, implying that the threshold criterion (3) leads to the same outcome for different competitors. This, however, cannot be the whole story, as we showed in the main text that random initialisation does lead to pronounced competitor assortment in a multi-patch scenario.

Our second reason highlights a difference between the two-patch scenario (which is the standard scenario considered in the literature) and a multi-patch scenario (as the one considered in our study). Threshold criterion (3) is based on inequality (2), which implicitly assumes that the decision-making individual compares two patches that it does not occupy. This is the case if individuals enter the system from the outside, and it is typically the case if many patches are compared with each other (as an individual can only occupy one of the patches, most patch comparisons involve patches not occupied by the individual). The situation is different in the two-patch scenario: if an individual makes a choice ‘from within’, it must already occupy one of the two patches under comparison. Let us call the occupied patch *P*_occ_ and the other patch *P*_other_. The individual should switch to the other patch if that other patch yields a higher intake rate:

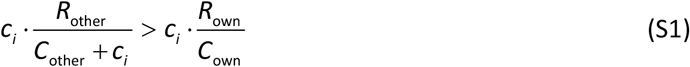

or, equivalently, if:

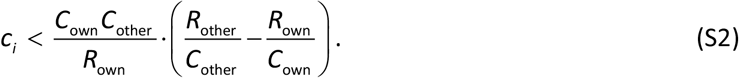

If the own patch has a higher resource availability (*R*_own_/*C*_own_ > *R*_other_/*C*_other_), the right-hand side of (S2) is negative, implying that individuals should never switch to the other patch, irrespective of their competitive ability. However, strong and weak competitors may differ in their patch preferences if the resource availability is higher on the other patch. Now, (S2) is a threshold criterion which is most likely satisfied for weaker competitors. This is in line with the findings of Houston & Lang (1998), who noticed that weak (but not strong) competitors may revise their earlier patch choice decisions once a strong competitor has moved into their patch. Notice that the 2-patch criterion (S2) does no longer contain the difference in resource richness (*R*_own_ − *R*_other_) in the denominator of the right-hand side. This means that the bias between strong and weak competitors is not based on differences in resource richness *per se*, but on differences in resource availability. Accordingly, one should not expect the assortment of strong competitors to resource-rich patches, in line with Figure S1 (blue line).

This is where the difference between a two-patch scenario and a multi-patch scenario becomes decisive. In a multi-patch scenario, relevant patch comparisons occur predominantly between patches not currently occupied, and therefore threshold (3) applies rather than (S2). Likewise, the increased number of patches makes diverging patch choice decisions between individuals of different competitive ability more likely. Extending the simulations of Figure S1 to multiple patches, we observe that some spatial assortment indeed occurs when individuals are initialized at random over ten patches instead of two (Figure S2, blue curves in the left panels), even if these patches are downscaled versions of patches A and B in the two-patch scenario. Previous theoretical treatments have predominantly focussed on the two-patch scenario, and this qualitative difference between two and multiple patches is therefore of some significance. We also observe a substantial increase of spatial assortment between two- and ten-patch scenarios if foragers are initialized outside of the patches (Fig. S2, red curves in the left panels).

By the same token, we can extend our simulations to consider the effect of more than two competitive types. Intuitively, the threshold criterion should become more relevant for a broader range of competitive types. Considering five instead of two competitive types, where competitive ability is given by *c*_*i*_ =*c*_1_/*i*, we observe strengthened spatial assortment for external initialization (Fig. S2, red curves in right panels). At random initialization (Fig. S2, blue curves in right panels) an increased number of types does not automatically lead to spatial assortment: On two patches, competitive types are distributed randomly independent of the number of types considered. Only when 10 patches are considered, does an increased number of types lead to some reinforcement of spatial assortment. Again, this is explained by the difference between equations (3) and (S2). For the simulations shown in Fig. S2, we used a population size of 2,000, but this parameter only affects the spread of the probability distributions and not their location.

**Figure S2:**
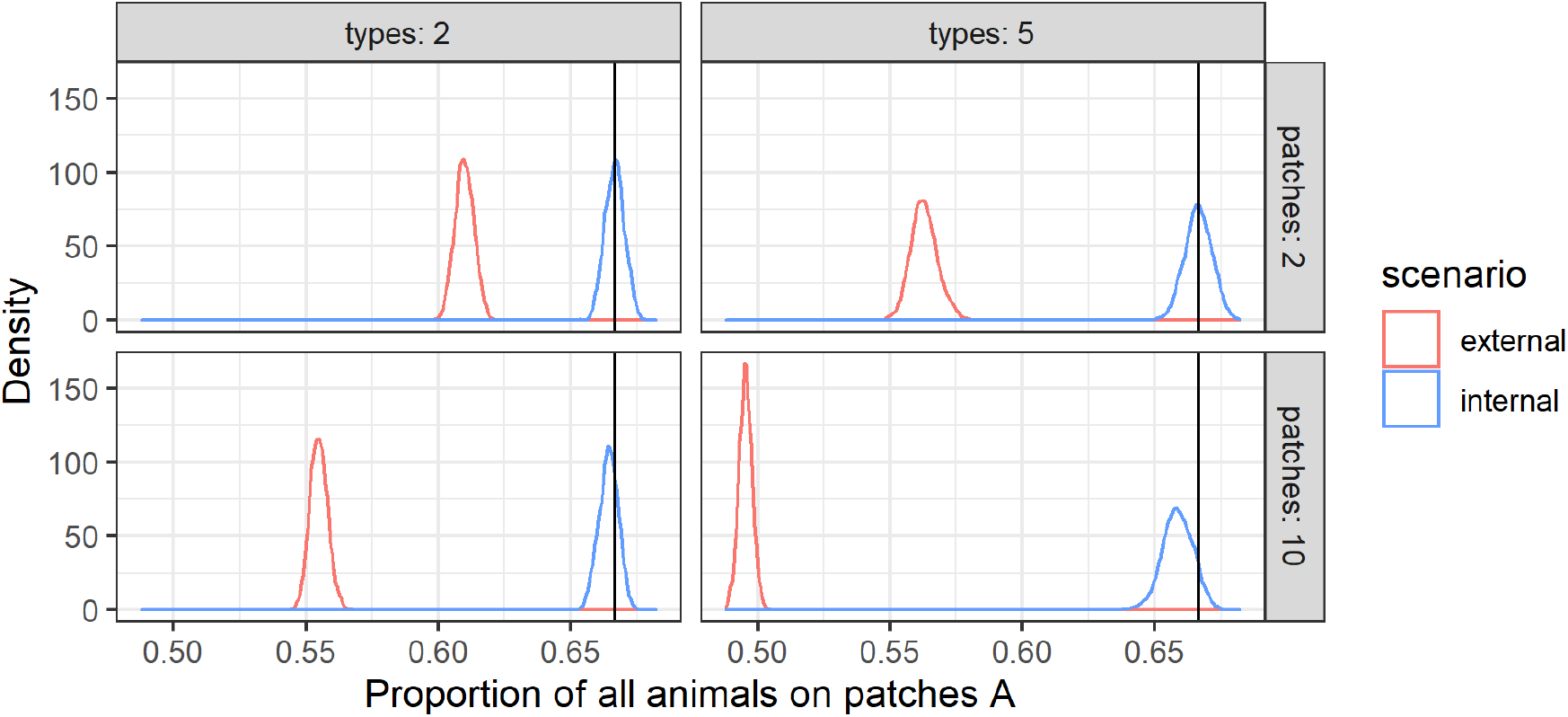
Effect of the number of patches and the number of competitive types on spatial assortment in two habitat choice scenarios. As in Fig. S1, the panels show the distribution of competitors over patches, based on 1,000 simulations for the external initialisation scenario (red) and the internal initialisation scenario (blue). The population now consists of 2,000 individuals, which can either be of two types (as in Fig. S1) or of five types, with competitive abilities *c*_*i*_ =*c*_1_/*i*. There are either two patches A and B (as in Fig. S1) or ten patches, where five are resource rich, while the other five are resource poor. As before, the resource influx in the resource-rich patches is twice as large as in the resource-poor patches.

